# Proteins with anti-apoptotic action in the hemolymph of caterpillars of the Megalopygidae family acts by maintaining the structure of the cellular cytoskeleton

**DOI:** 10.1101/2023.02.10.527989

**Authors:** Nathalia Delazeri de Carvalho, Henrique Krambeck Rofatto, Karina de Senna Villar, Roberta Fiusa Magnelli, PI da Silva Junior, Ronaldo Zucatelli Mendonça

## Abstract

Brazil has a very large biological variety, which is an almost inexhaustible source of substances of pharmacological and biotechnological interest. Several studies have demonstrated the presence of bioactive peptides in insect hemolymph and their potential use as therapeutic agents. However, few data are available regarding molecules extracted from insects with anti-apoptotic action. The objective of this work was to identify and isolate proteins from the hemolymph of caterpillars of the *Megalopygidae* family with pharmacological and biotechnological interest. Two species of this family were studied, *Podalia sp and Megalopyge albicolis*. Cytotoxicity tests on Vero and Sf-9 cells revealed that the hemolymph of both caterpillars was cytotoxic only at concentrations greater than 5%v/v. In the anti-apoptotic activity assays, it was verified that the supplementation of cell cultures with only 1% of hemolymph v/v is sufficient to inhibit cell death by apoptosis induced by different inducers such as terbutyl, actinomycin D, hydrogen peroxide, or even by nutrient depletion. For this study, cells were stained with trypan blue, crystal violet, and fluorescent markers to cytoskeleton (actin and tubulin), mitochondria membrane electric potential (JC-1), and apoptosis marker (acridine orange and ethidium). The protein responsible for anti-apoptotic action was isolated through gel filtration chromatography, using an AKTA purifier high-resolution liquid chromatography system. The hemolymph was fractionated into 3 pools for *Podalia sp* and 6 pools for *M. abicolis*. In the antiapoptotic tests, semi-purified hemolymph from both caterpillars showed anti-apoptotic effect in VERO and SF-9 cells, pre-treated with only 1% v/v of hemolymph and induced to death by different and apoptotic inductors. Was observed that the molecule with anti-apoptotic effect is present in pool 3 in both hemolymphs. This protector effect blocked and attenuated the disruption of the cytoskeleton (actin filaments), being that the protective effect also was observed on the integrity of the mitochondrial membrane of SF-9 cells pre-treated with both hemolymphs and treated with the apoptosis inducer Terbutil at concentrations of 25 to 100μM. By acting on the mitochondrial pathway of death by apoptosis, a pathway that can cause disorders and diseases neurodegenerative such as Parkinson’s and Alzheimer’s diseases, substances present in the hemolymph of these and others caterpillars could be good candidates in studies for the treatment of these diseases.

## INTRODUCTION

Insects are believed to have emerged in the Devonian period, over 350 million years ago. During this period, insects lived and survived in almost all ecological niches on the planet, from deserts, polar regions, contaminated environments,, or extremely harmful to the development of life, developing protection and defense mechanisms for each adverse situation. Several studies have been carried out demonstrating the presence of pharmacologically active substances in the hemolymph of insects with antiviral action (Mendonça et al, 2023; Carvalho et al, 2022, Carmona et al, 2020; Coelho et al, 2018, Greco et al, 2009, Carmo et al, 2012), antimicrobial and antifungal action (Cantero et al, 2022, Hayashida et al 2019, Zhu et al., 2000, Gross et al., 1998; Johns et al., 1998; Lanz-Mendoza et al., 1996; Lowenberger et al., 1999) antifreeze (Kristiansen et al, 2005) hormonal (Huberman et al., 2000; Jones et al., 1993; Kurata et al., 1994; Lin et al., 1998), enzymatic effects, among others. (Guerrero et al., 1999; Hamdaoui et al., 1998; Jiang et al., 1999; Rosenfeld and Vanderberg, 1998; Shiotsuki et al., 2000; Yamamoto et al., 1999). Some works have been published demonstrating the positive effect of hemolymph in promoting growth (Kanaya and Kobayashi, 2000; Rhee et al., 1999; Rhee and Park, 2000), production of recombinant proteins (Vieira et al, 2010, Mendonca et al., 2008; Woo et al., 1997). Kanaya and Kobayashi (2000) isolated and characterized a protein present in the hemolymph of the silkworm (Bombyx mori) capable of increasing the activity of a recombinant protein (luciferase) by approximately 10,000 times. Some studies have been carried out seeking to isolate and characterize the factors involved in these effects (Nussbaumer et al., 2000; Ochanda et al., 1992; Shishikura et al., 1996; Shishikura et al., 1997) however, only some studies have demonstrated the site of action of these proteins. We have verified the occurrence of cell death by apoptosis in cell cultures depleted of nutrients (Mendonca et al., 2002; Meneses-Acosta et al., 2001) or by induction of chemical agents (Souza et al., 2005). We have shown that supplementation of insect and mammalian cell cultures with *Lonomia obliqua* hemolymph can extend culture viability by preventing death by apoptosis (Maranga et al., 2003; Raffoul et al., 2005; Souza et al., 2005). Recently we have observed that this antiapoptotic effect of hemolymph is associated with the mitochondria pathway, acting in the mitochondria membrane permeabilization (MMP) and with the maintenance of the electrochemical potential of the mitochondria (Mendonça and Martins, 2022). These information makes hemolymph an interesting source of additives for cell cultures. However, studies with caterpillars of the *Megalopygidae* family are very scarce. Megalopygids (hairy caterpillars) are generally solitary, and non-aggressive, from 1 cm to 8 cm in length. Its dorsal “hairs” are long bristles, fragile, silky, and harmless, of varied colors that camouflage the true pointed and stinging bristles. The base of the bristle is a portion implanted in the caterpillar’s skin, consisting of a single gland, filled with a toxic substance. When pressed upon contact, the gland releases the poison that travels through a channel, being injected into the human skin and causing injury. Some species of caterpillars are especially beautiful and attractive, which makes people, especially children, run their hands over the caterpillar. Species of Megalopygids can vary in color from gray to red, pink, yellow, black brown.

Some popular names are given to these caterpillars such as caterpillar-dog, flannel caterpillar, caterpillar-kitten, hat-armor, and bug-hairy.

In Brazil, there are approximately 50,000 species of lepidopterans, about 220 species in the Megalopygidae family. These caterpillars are found mostly in the neotropics regions (Cardoso and Junior 2005). Megalopygidae and the Saturniidae family *(Lonomia obliqua)*, are responsible for the majority of accidents by caterpillars in Brazil. Reports of irritating properties of lepidopterans are described by Pliny the Elder and Galen during the Roman Empire (Fonseca, F, 1950). The first reports in the American continent were made by priest José de Anchieta in his Letters from São Vicente” (1569) and Marcgrave and Piso, the fathers of Natural History in Brazil, recorded cases of accidents with caterpillars in the Northeastern region, in 1658 (Rotberg A, 1990). Megalopygid venom can cause an extreme allergic reaction, resulting in rashes, inflammation, blisters, and breathing difficulties. The principal species in this family are *Podalia sp* and *Megalopyge albicolis*.

Studies aiming at the search for substances with pharmacological or biotechnological action in Megalopygidae family are practically non-existent. We recently carried out a study with these two caterpillars to identify substances with antiviral action. In this work we identified substances with potent antiviral action against the Measles virus, Influenza virus (H1N1), Herpes Simplex Virus and picornavirus (Carvalho et al, 2022). Studies on cell death are of great importance. When the cell dies, it guarantees the regulation of the body’s homeostasis and important functions, such as organ remodeling and control of the immunity system. The entire process of controlling cell death, known as apoptosis, is regulated by a network of proteins. Knowledge of cell death mechanisms can lead to important strategies in the control of various physiological processes. We previously identified substances with potent anti-apoptotic action in the hemolymph of caterpillars of the Saturnidade family (Souza et al., 2005) and *Megalopyge* (Mazzoni et al, 2014). So, due to the importance of studies on cell death, the objective of this research is the identification, isolation, and purification of substances with anti-apoptotic action in the hemolymph of caterpillars of the Megalopygidae family.

## 3. MATERIALS AND METHODS

### 3.1 Cell culture

#### 3.1.1 Cell line, culture medium and cell culture

##### VERO cells

(Cercophitecus aethiops African green monkey kidney fibroblast cells), were cultured at 37ºC in T-flasks containing 5 ml of Leibovitz medium (L-15) supplemented with 10% fetal bovine serum. Cell cultures were also performed in 6, 12, 24, or 96 well microplates.

##### Sf-9 cells

(Spodoptera frugiperda pupal ovary strain) were cultured at 29°C and 120 rpm in Schott flasks with a working volume of 13 mL in SF-900 medium or microplates.

Cell growth and morphology were monitored using an Olympus CK2 inverted optical microscope at 100, 200, or 400x magnification.

### 3.2 Isolation and purification of proteins

#### 3.2.1 Hemolymph

The hemolymph used in this study was obtained from caterpillars of the Megalopygidae family (*Podalia sp and M. albicolis*), by hemolymph leakage after cutting the caterpillar’s pseudo feet. The hemolymph thus collected was stored at –20°C. Before use, the hemolymph was thawed, centrifuged at 1,000 g for 10 minutes, and filtered through a 0.22 μm sterilizing membrane.

The material was kept at −4°C and used as a supplement to the cultures at a concentration of 1% to 2% (v/v).

#### 3.2.2 Hemolymph fractionation by chromatography

For the fractionation of *Podalia sp and M. albicolis* hemolymph, a high-resolution liquid chromatography system AKTA purifier (Amersham Pharmacia Biotech) was used. Its consistis of two pumps, ultraviolet light reader, programmer – Unicorn 4.0 software, and fraction collector Frac 900 (Amersham Pharmacia Biotech) coupled. 0.5 mL sample of hemolymph was subjected to gel filtration chromatography on “Superdex 75” columns in Tris buffer (20mM) and Nacl (150mM) with a flow rate of 0.5 mL/min. The fractions obtained were pooled.

#### 3.2.3 Determination of protein concentration

The protein concentration present in the hemolymph and in its fractions was determined by the NanoDrop 2000. The absorbance reading was performed at 280 nm.

#### 3.2.4 SDS-PAGE polyacrylamide gel electrophoresis

Electrophoresis of polyacrylamide gels, for visualization of the protein bands of the total hemolymph and its fractions, was carried out according to (Laemmli, 1970), under denaturing conditions, in the presence of sodium dodecyl sulfate (SDS).

### 3.3 Cytotoxicity e Genotoxicity of hemolymph

#### 3.3.1 Cytotoxicity of hemolymph

To evaluate cytotoxicity, 0.5; 1; 2, or 5% of total hemolymph or its fractions obtained by chromatography was applied to cell culture. The morphology of the cultures was monitored through daily observation in an inverted optical microscope at 100x and 200x magnification. At the end of the cultivation, the wells were stained with crystal violet (0.25%) and photographed, determining the morphological changes that occurred in the cells.

The cytotoxicity on Vero cells of both hemolymph by apoptosis inducers cells was also evaluated by the MTT method, as described by Mosmann, 1983). For this, Vero cells were treated with 2% v/v of hemolymph or tert-butyl at concentrations 0,5, and 1mM for 18 hours. The medium was removed and 100μl of a 0.5 mg/ml MTT solution was added and the plate was again incubated at 37°C for 4 hours. Next, the supernatant was removed and 100μl of DMSO was added. The formazan crystals were then solubilized by hand stirring. Absorbance was determined in a microplate reader at 540nm. The ability of cells to reduce MTT provides an indication of mitochondrial activity and integrity, which are interpreted as measures of cell viability. The technique was first described to evaluate compounds with antitumor activity, which is closely related to cytotoxicity.

#### 3.3.2 Genotoxicity of hemolymph

The genotoxicity of raw hemolymphs and its fractions obtained by gel filtration chromatography was evaluated by the comet method as described by Magneli et al, (2021).

### 3.4 Chemical inducers of apoptosis

Tert-butyl hydrogen peroxide (t-BPH) (25 to 2000 μM), Hydrogen Peroxide (H2O2) (25 to 2000 μM), and Actinomycin D (200 and 800ng/mL) were used as apoptosis chemical inducers.

### 3.5 Characterization of the biological activity of hemolymphs and its fractions

#### 3.5.1 Effect of *Podalia sp* hemolymph in promoting cell growth

The total hemolymph of ***Podalia sp*** was tested in **Sf-9 cell** cultures, seeking to identify its ability to promote cell growth. For this, 1% of the total hemolymph v/v was added, at the time of cell inoculum to the culture of Sf-9 cells. Culture samples were obtained daily and the number of viable cells was determined by the trypan blue staining method.

#### 3.5.2 Anti-apoptotic action of hemolymphs

##### 3.5.2.1 Protector effect of *Podalia sp* hemolymph in death induced by nutrient depletion

The effect of total hemolymph and its fractions on **VERO cell** growth in culture with a depleted medium was analyzed. To this, after the cell’s confluence, the medium culture was changed by PBS. At the time of nutrient depletion, 1% v/v of total hemolymph or its fraction 3 obtained from gel filtration column chromatography was added to the and analyzed for 96 hours.

##### 3.5.2.2 Protector effect of Podalia sp and M. albicolis hemolymph in death induced by t-BPH and H_2_O_2_

The apoptotic effects of Podalia sp. and M. albicolis hemolymph protect against cell death induced by chemical inducers of apoptosis. VERO cells were pre-treated with 1 or 2% (v/v) crude hemolymph for 1 h. After this period, cell death was induced with 0.5mM or 1 mM t-BPH or H2O2 (25, 50, 100, or 400μM), for 18 h. The cells were stained with crystal violet and observed under an inverted optical microscope (400x).

#### 3.5.3 Determination of cell death by epifluorescence microscopy, confocal microscopy and optical microscopy

The total hemolymph of *Podalia sp*. or *M. albicolis* and its fractions, obtained by chromatography, were tested in cell cultures, seeking to verify factors present in the hemolymph that promote cellular protection against death by apoptosis. Total hemolymph (1% v/v) or its fractions (2% v/v) were added to the cultures of Vero and Sf-9 cells, and after 1 h, cell death inducers were added. After 2 days, for Vero cells or 5 days for Sf-9 cells, after cell contact with the inducer, the cultures were stained and analyzed using epifluorescence optical microscopy. To this, 5 μL of a solution containing 100 μg/ml acridine orange (AO) and 100 μg/ml ethidium bromide (EB) was added to 100 μL of cell culture (Vero or Sf-9) at a concentration between 0.5 and 1 × 106 cells/mL. This mixture was examined using epifluorescence microscopy with ultraviolet light, a blue filter, and confocal microscopy. Apoptosis was also visualized by light microscopy after staining with trypan blue.

#### 3.5.4 Determination of cell death by flow cytometry

Vero cells were treated or not with total hemolymph from *Podalia sp*. Or *M. albicolis* (1% v/v) and after 1-hour t-BPH or Actinomycin D was added. After 18 h, samples containing 106 cells were obtained. These were centrifuged at 800 g for 5 min, and the pellet was fixed in ice-cold 70% ethanol and stored at −20°C. On the day of analysis, ethanol was removed by centrifugation (800 g for 5 min) and cells were incubated with 1 mL of Vindelev lysis buffer (1 mg/ml citrate, 50 μg/mL propidium iodide, 1 mL NP −40 (0.1%), 700 U/ml RNase, 0.01 M NaCl). After 10 min, the cells were centrifuged (10000 g for 5 min), and the pellet was resuspended in 1 mL of FACS buffer. The samples were processed in a FACS Calibur (BD Biosciences) with a wavelength of 488 nm for excitation and 620 nm for emission to determine the different phases of the cell cycle.

#### 3.5.5 Determination of alteration of the cytoskeleton of Vero cells by phalloidin-FITC labeling

One of the first changes observed in apoptotic cells was the loss of cell adhesion due to alterations in the cytoskeleton caused by microfilament retraction. To visualize actin filaments, Vero cells were labeled with FITC - conjugated phalloidin. Nuclei were co-labeled with propidium iodide. The cells were seeded on plates with coverslips at a maximum density of 105 cells/mL and after 24 hours they were pretreated or not with the total hemolymph of *Podalia sp* or *M. albicolis* (1% v/v) for 1 h and then induced with different doses of tert-butyl (25, 50 and 100μM) for 4 h. The cell medium was then removed and the cells were fixed with 4% paraformaldehyde in PBS for 20 min at room temperature. Cells were washed 3 times/5 minutes with PBS. The cells were then permeabilized with 0.1% Triton X-100 in PBS at room temperature for 15 min. After this time, a block was performed with PBS+BSA + 1% BSA + RNase (100μg/mL), for 40 min at 37 °C. These cells were then incubated with FITC - conjugated phalloidin (20μg/ml) diluted in PBS+BSA + 1% BSA for 3 h at 37°C. Cells were washed 3 times for 1 min each with PBS, and incubated with propidium iodide (20 μg/ml) diluted in PBS+BS + 1%, for 1.5 h at 37°C. The cells were again washed 3 times/5 minutes with PBS. The slides were then observed under a confocal microscope (LSM 510 META, ZEISS). Labeling with phalloidin-FITC was followed by laser starting at 488 nm with an emission filter BP500-550 nm, while for iodide, it was used to study the laser at 543 nm with the emission filter LP560 nm.

#### 3.5.6 Labeling of tubulin in Vero cells

To visualize tubulin, Vero cells were labeled with anti-tubulin antibodies. Additionally, the nuclei were co-labeled with propidium iodide. The cells were seeded in microplates containing coverslips at a maximum density of 105 cells/mL. After 24 h, the cell medium was removed and the cells were fixed with 4% paraformaldehyde in PBS for 20 min at 37°C. Cells were washed 3 times/5 minutes with PBS. The cells were then permeabilized with 0.1% Triton X-100 in PBS at room temperature for 10 min. Next, blocking was performed with PBS+ Triton X-100 0.05% + FBS 5% + RNAse (100μg/mL), for 40 min at 37 °C. Cells were washed three times for 1 min each with PBS and 5% FBS. Cells were again washed 3 times/5 minutes with PBS. Cells were incubated with the primary antibody, anti-beta-tubulin produced in mice (Sigma), (1 μg/mL + PBS + 0.05% Triton X-100 + 5% FBS) at 4°C overnight. The cells were washed 3 times/5 minutes each with PBS. Then, the cells were incubated with blocking solution for 2 h at room temperature with the secondary antibody, anti-mouse IgG conjugated to Alexa Fluor 488 (Life Technologies) (10 μg/mL + PBS + Triton X-100 0.05% + 5% FBS at room temperature for 2 h). The cells were then washed for 2 times/5 minutes each with PBS. Propidium iodide (10μg/mL) was added for 1 h, after which the cells were again washed 3 times/5 minutes each with PBS. The slides were then mounted with VectaShield and sealed with enamel. Slides were analyzed under a confocal microscope (LSM 510 META, ZEISS). Tubulin labeling was analyzed with a 488 nm laser for excitation and with a BP500-550 nm emission filter, while for iodide a 543 nm laser was used for excitation with an LP560 nm emission filter.

#### 3.5.7 Determination of mitochondrial membrane potential (ΔYm)

For the determination of the effect of hemolymph in protecting tert-butyl-induced apoptosis (t-BPH), Sf-9 cells were cultured in 12-well microplates. After reaching semi-confluence, the cells were treated with 1% (v/v) crude hemolymph. After 1 h of contact, cells were treated with 600 μM t-BPH. The cultures were kept in an oven for 4 h, after which the cells were washed with PBS and treated with the dye JC-1 (5,5’,6,6’-tetrachloro-1,1’,3,3’-tetraethylbenzimidazolylcarbocyanine) (2μM). The cells were analyzed and photographed using a confocal microscope.

## 4. RESULTS AND DISCUSSION

### 4.1 Determination of cell death by epifluorescence microscopy, and optical microscopy

Apoptosis in **Vero** and **Sf-9** cells was observed by epifluorescence microscopy with ultraviolet light and blue filter after being stained with acridine orange and ethidium bromide **(figure 1)**. Apoptosis was visualized by light microscopy after staining with trypan blue, as shown in **figure 2**. The images in Figures 1 and 2 are representative of all the experiments carried out to determine apoptosis by microscopy.

**Figure 1:**
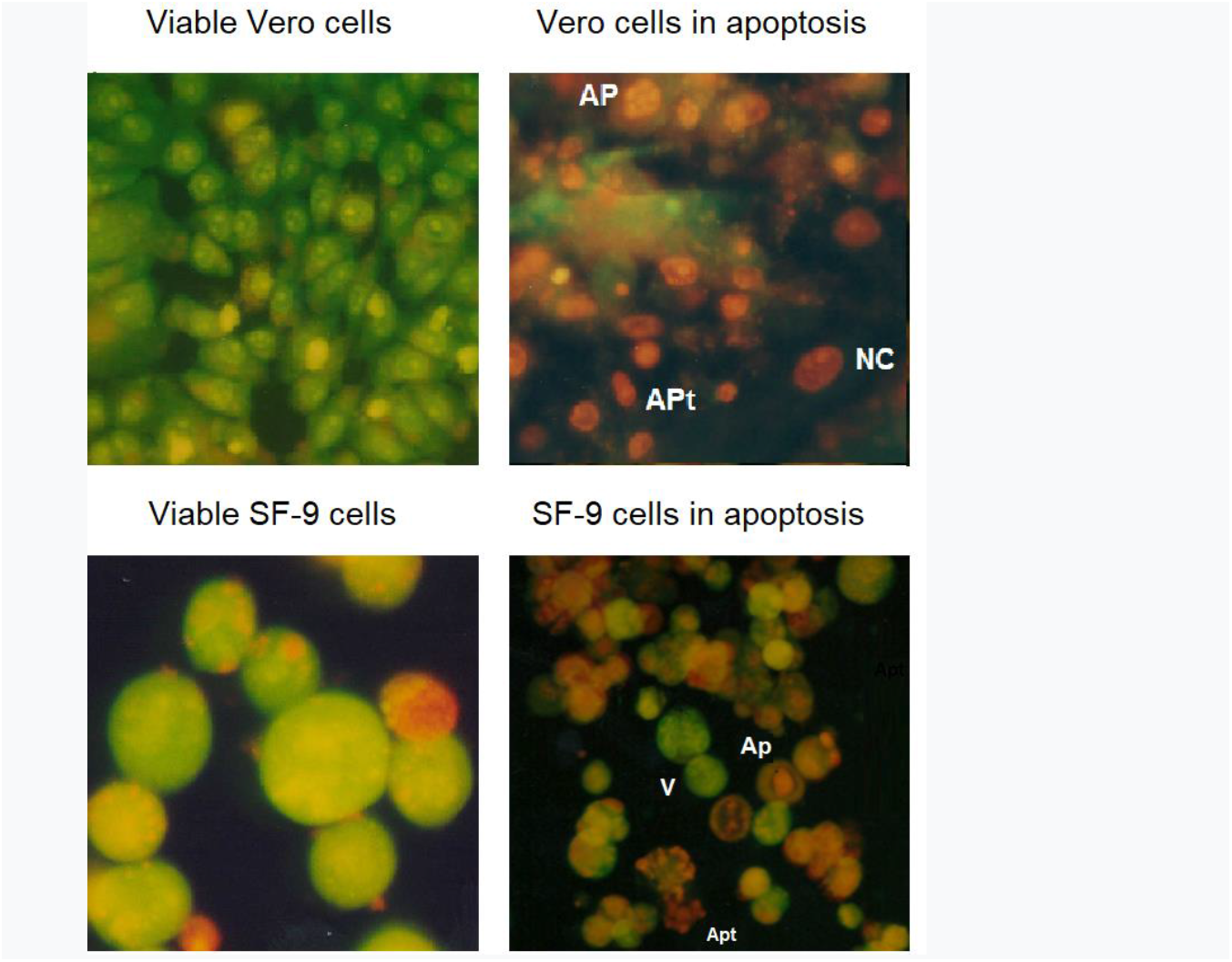
Photomicrographs of Vero and Sf-9 cells. Cells at different stages of cell death were stained with acridine orange and ethidium bromide. Viable cells (V) are indicated in green. Cells in red are observed in the process of early apoptosis (AP) or late apoptosis (APt). In addition, the cells were observed during the necrosis process (NC). Cells were observed under an epifluorescence microscope (100x (or 400x)

**Figure 2:**
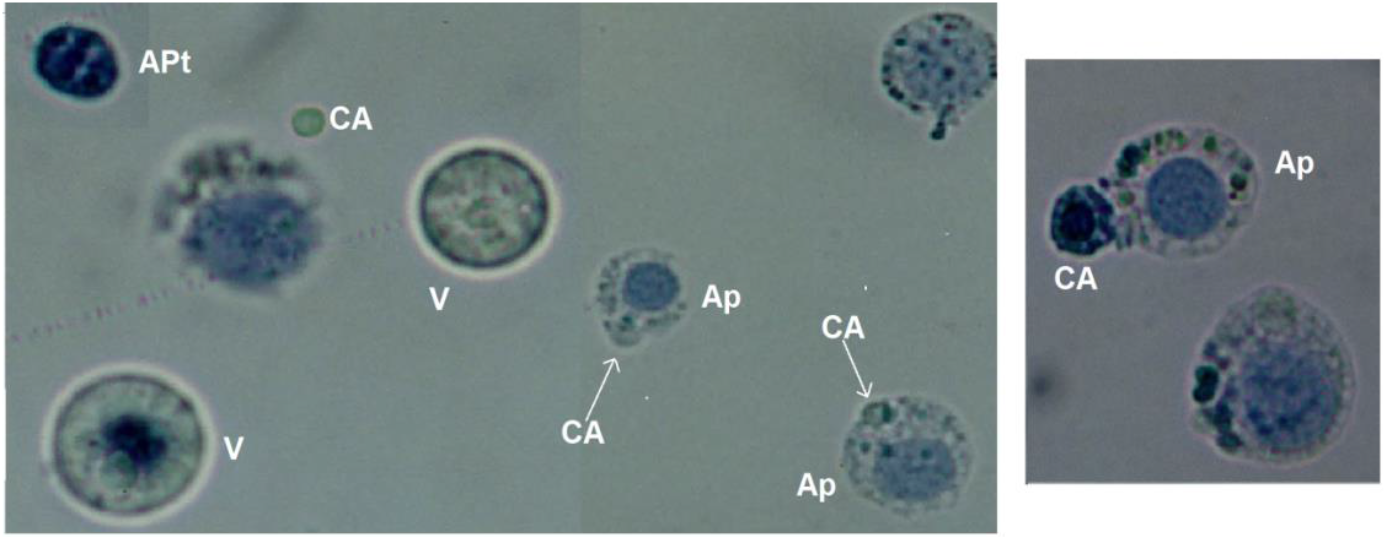
Photomicrograph of Sf-9 cells showing viability (v) or at different stages of apoptosis (AP). The formation of apoptotic bodies (CA) and cells with later apoptosis (APt) were also visible. A sample of the Sf-9 cell culture was obtained on day 5, stained with trypan blue, and visualized under an optical microscope. (400x).

### 4.2 Hemolymph fractionation by chromatography

To determine which fraction was responsible for the observed effects, total hemolymph from Podalia sp. and M. albicolis was semi-purified in a gel filtration column (Superdex 75). The obtained fractions were pooled into three pools for Podalia sp. and six pools for M. albicolis and tested for their anti-apoptotic activity. In the first step of purification in gel filtration columns, the molecular weight profile of the different proteins that make up the hemolymph was determined. The chromatographic profile is shown in **figures 3 and 4** and the polyacrylamide gel of fractions in **figure 5**

**Figure 3:**
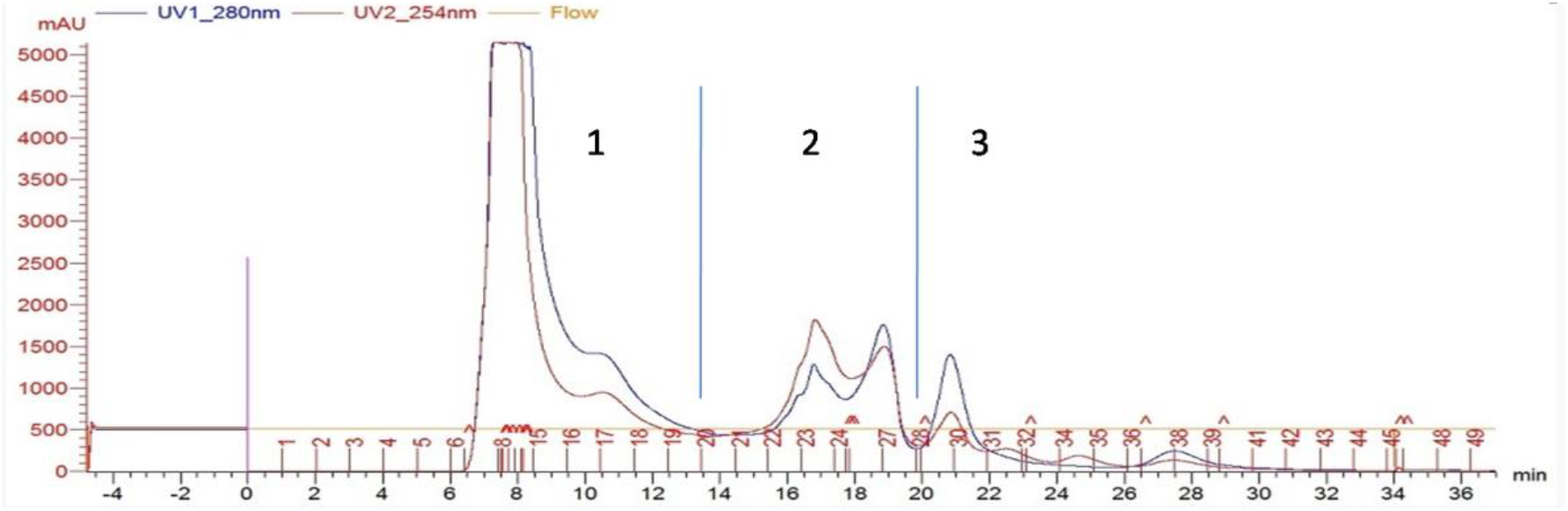
Purification of Podalia sp. hemolymph by gel filtration chromatography on a “Superdex 75” column. Elution was performed using Tris (20mM) and Nacl (150mM) solution. The flow used was 0.5 ml/minute and fractions of 1 ml were collected. The fractions were separated into 3 fractions as indicated by the bars

**Figure 4:**
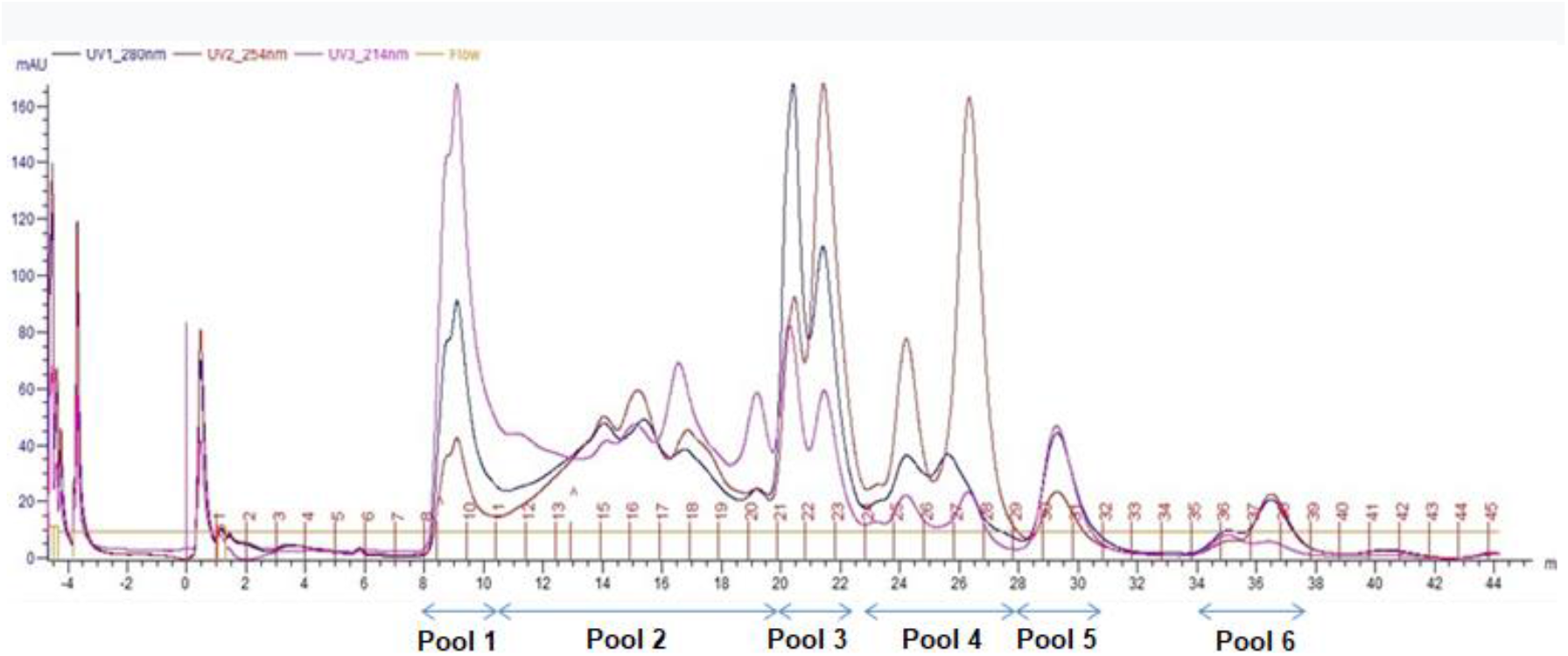
Purification of ***M. albicolis*** hemolymph by gel filtration chromatography on a “Superdex 75” column. Elution was performed with Tris (20mM) and Nacl (150mM) solution. The flow used was 0.5 ml/minute and fractions of 1 ml were collected. Fractions were separated into 6 groups as indicated

**Figure 5:**
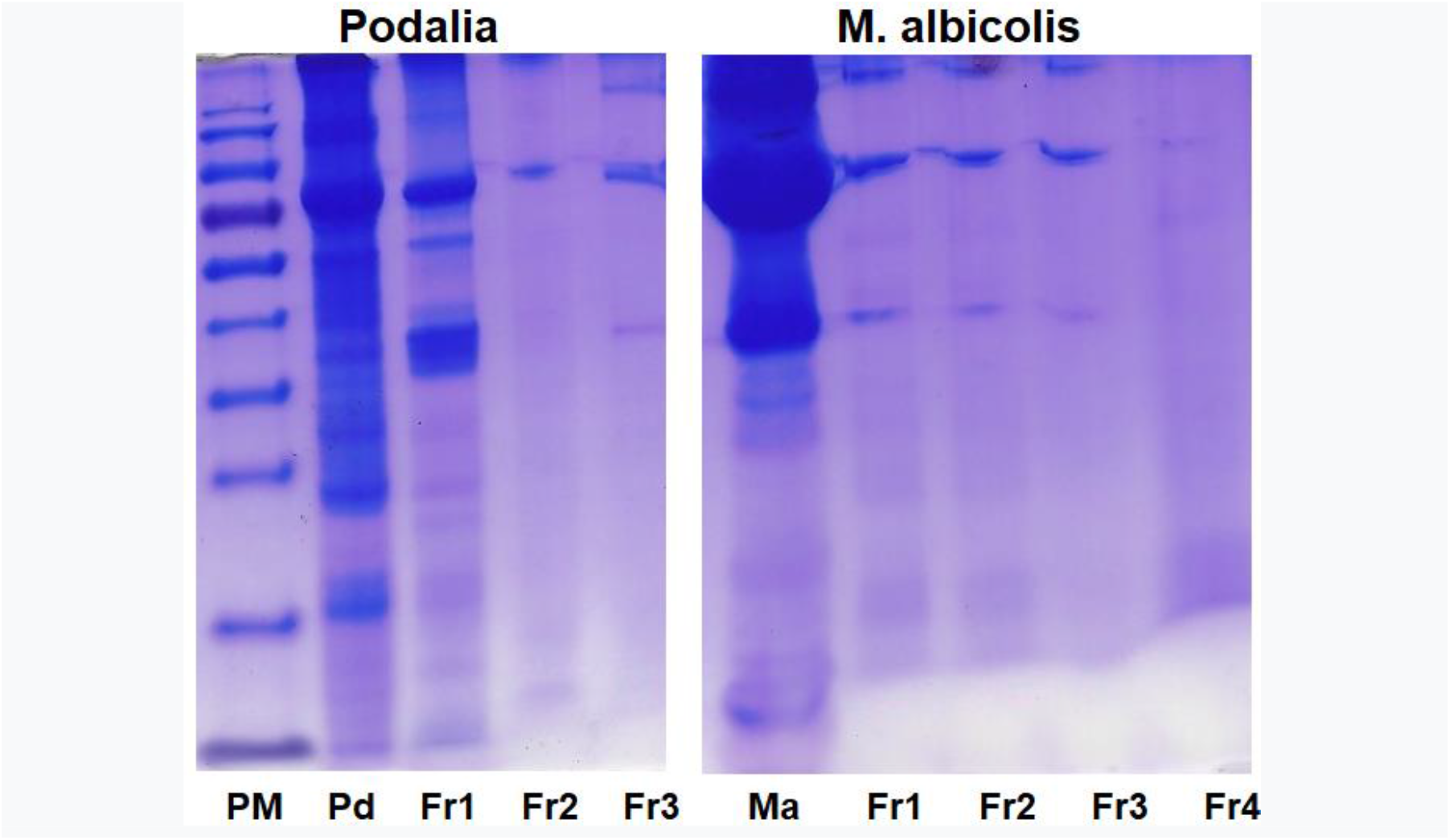
12% SDS-PAGE gel electrophoresis. Crude hemolymph from Podalia sp. (Pd), M. albicolis, and its fractions were obtained by gel filtration chromatography, and column 1 (PM) corresponded to the calibration molecular weight. Pd (Podalia sp crude). Ma (crude M. albicolis). The gel was stained with Coomassie Blue dye.

#### 4.2.1 Determination of protein concentration

The protein concentration present in the hemolymph and its fractions obtained by gel filtration chromatography was determined using a NanoDrop 2000. Absorbance was measured at 280 nm. The results are presented in **Table 1**.

**Table 1.** Measurement of protein concentration in the crude hemolymph of Podalia sp. and M. albicolis and the fractions obtained from purification of the hemolymph by gel filtration chromatography on a “Superdex 75” column.

| *Podalia* SP | mg/mL | *M. albicolis* | mg/mL |
| --- | --- | --- | --- |
| Hemolymph crude | 41,012 | Hemolymph crude | 42,591 |
| Pool 1 | 1,060 | Pool 1 | 1,085 |
| Pool 2 | 1,943 | Pool 2 | 0,259 |
| Pool 3 | 0,828 | Pool 3 | 0,383 |
|  |  | Pool 4 | 0,046 |
|  |  | Pool 5 | 0,129 |
|  |  | Pool 6 | 0,100 |

#### 4.2.2 Cytotoxicity

The cytotoxicity of the total hemolymph of *Podalia sp*. and *M. albicolis* was determined in **Vero** and **Sf-9** at concentrations of 0.5, 1, 2, 5, and 10% v/v. Hemolymphs were cytotoxic only at concentrations greater than 5% of the final culture volume, demonstrating that their application in smaller amounts in cultures is feasible. At lower concentrations, no significant cell death was observed for any of the tested cells. Based on these results, 1% and 2 % (v/v) were used in all tests (data not shown).

#### 4.2.3 Genotoxicity of *Podalia sp* and *M. Albicolis* hemolymph

The genotoxicity of crude hemolymph from *Podalia sp*., *M. Albicolis*, and its fractions obtained by gel filtration chromatography was evaluated using the comet method. In our study, 90% of the comets were classified as class 0 (not damaged) or class 1 (little damage), indicating that the tested material was safe and non-genotoxic (data not shown).

### 4.3 Effect of Podalia sp hemolymph in promoting cell growth

The effect of total hemolymph on **SF-9** cell growth was also analyzed.

As shown in **figure 6**, the culture treated with hemolymph did not affect the number of cells; however, we observed a prolongation of cell viability, an indication of protective action.

**Figure 6:**
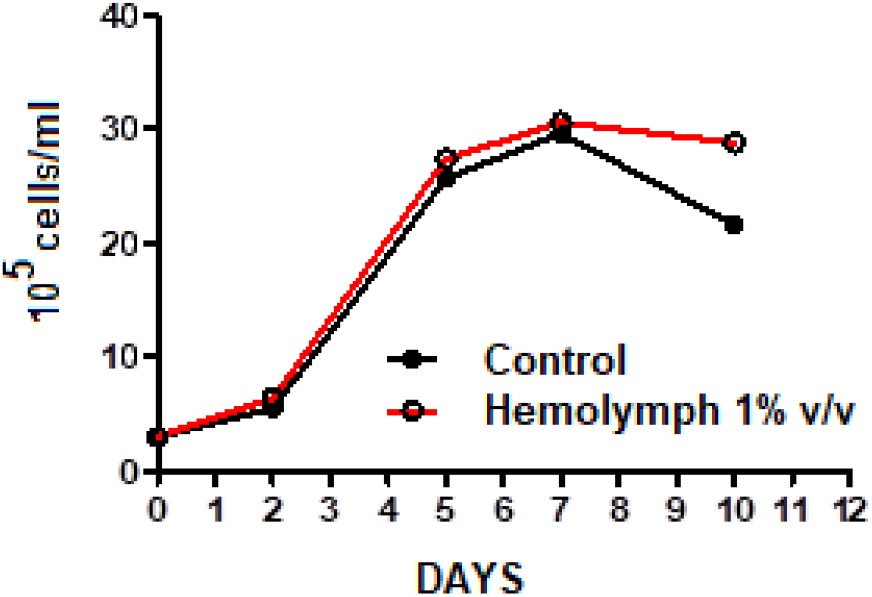
Cultures of SF-9 cells in Schott flasks containing 13 ml of Sf-900 medium maintained at 29°C and 120 rpm. Cultivation was initiated at a density of 3 × 105 cells/ml. Samples were obtained daily and the number of viable cells was determined by trypan blue staining.

#### 4.3.1 Protector effect of Podalia sp hemolymph in death induced by nutrient depletion

The effect of total hemolymph and its fractions on **VERO** cell growth in cultures with depleted medium was analyzed. To this end, the medium culture was changed with PBS and analyzed for 96 h. As shown in **figure 7**, nutrient withdrawal leads to cell death. However, when *Podalia sp*. hemolymph or it **fraction 3** was obtained using gel filtration chromatography, the morphology of the culture was maintained.

**Figure 7:**
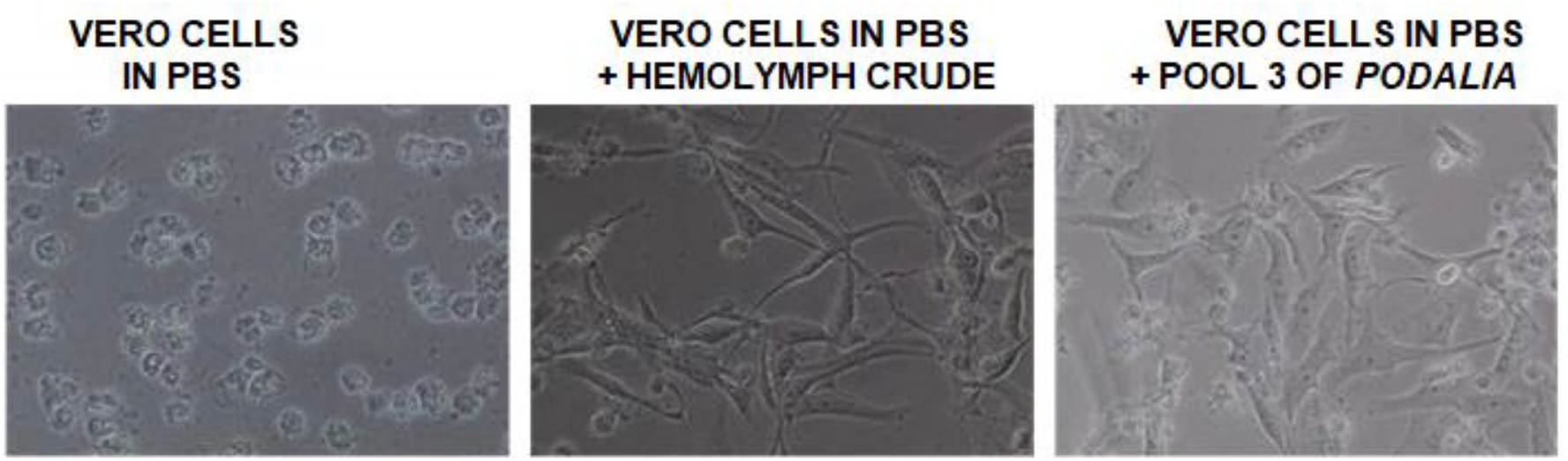
Effect of ***Podalia sp*** hemolymph on protection from cell death induced by nutrient depletion. At the time of nutrient depletion, 1% v/v of total hemolymph or its fraction 3 obtained from gel filtration column chromatography was added to the culture. After 96 hours the cells were observed in an inverted optical microscope (400x).

We also tested the ability of ***M. albicolis*** hemolymph (total) to protect against apoptosis induced by nutrient depletion. To this end, **VERO** cells were maintained in PBS for 96 h with 1% v/v total M. albicolis hemolymph. As shown in **Figure 8**, the removal of nutrients leads to cell death. However, when *M. albicolis* hemolymph was added to the culture depleted of culture medium, the culture morphology was maintained

**Figure 8:**
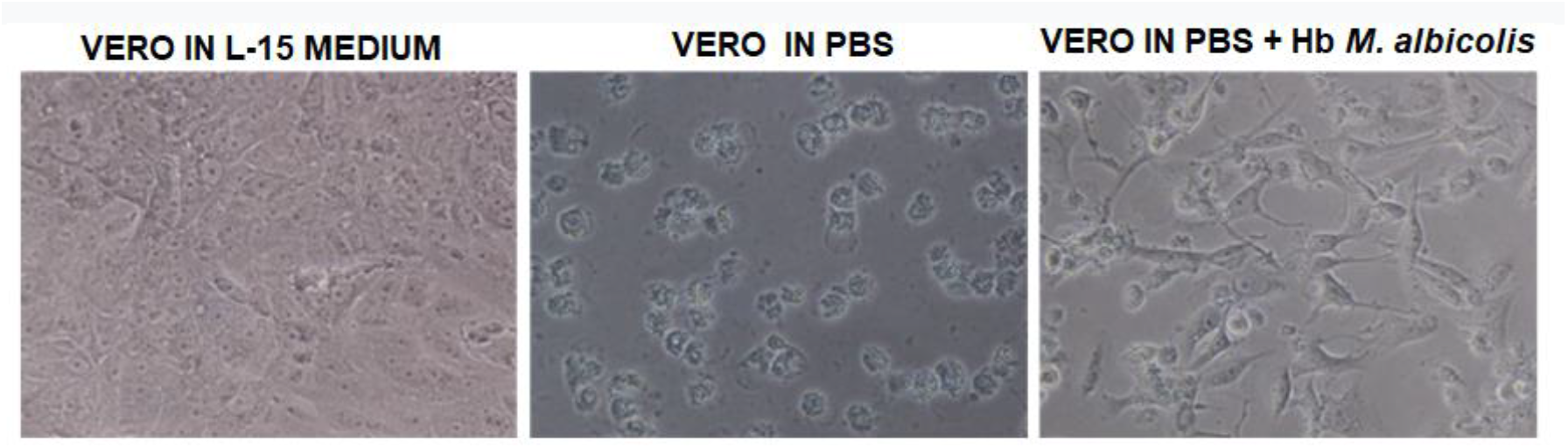
Effect of *M. albicolis* hemolymph on protection from cell death induced by nutrient depletion. At the time of nutrient depletion, 1% v/v total hemolymph was added to the culture. As a control, wells from the same culture plate were maintained in L15 medium containing 10% fetal bovine serum. After 96 h, cells were observed under an inverted optical microscope (400x).

### 4.4 Effect of apoptosis inducers on Sf-9

The action of chemical inducers of apoptosis, t-BPH (2mM) Actinomycin D (200 ng/ml), and Hydrogen Peroxide (H_2_O_2_) (2mM), was analyzed in **Sf-9** cells. The cultures were kept in contact with the inducer for 18 h and cell viability was determined using trypan blue. As shown in **figure 9**, all three inducers were able to induce cell death at the tested concentrations, 200ng/ml for actinomycin and 2 mM for t-BPH 2 H_2_O_2_.

**Figure 9:**
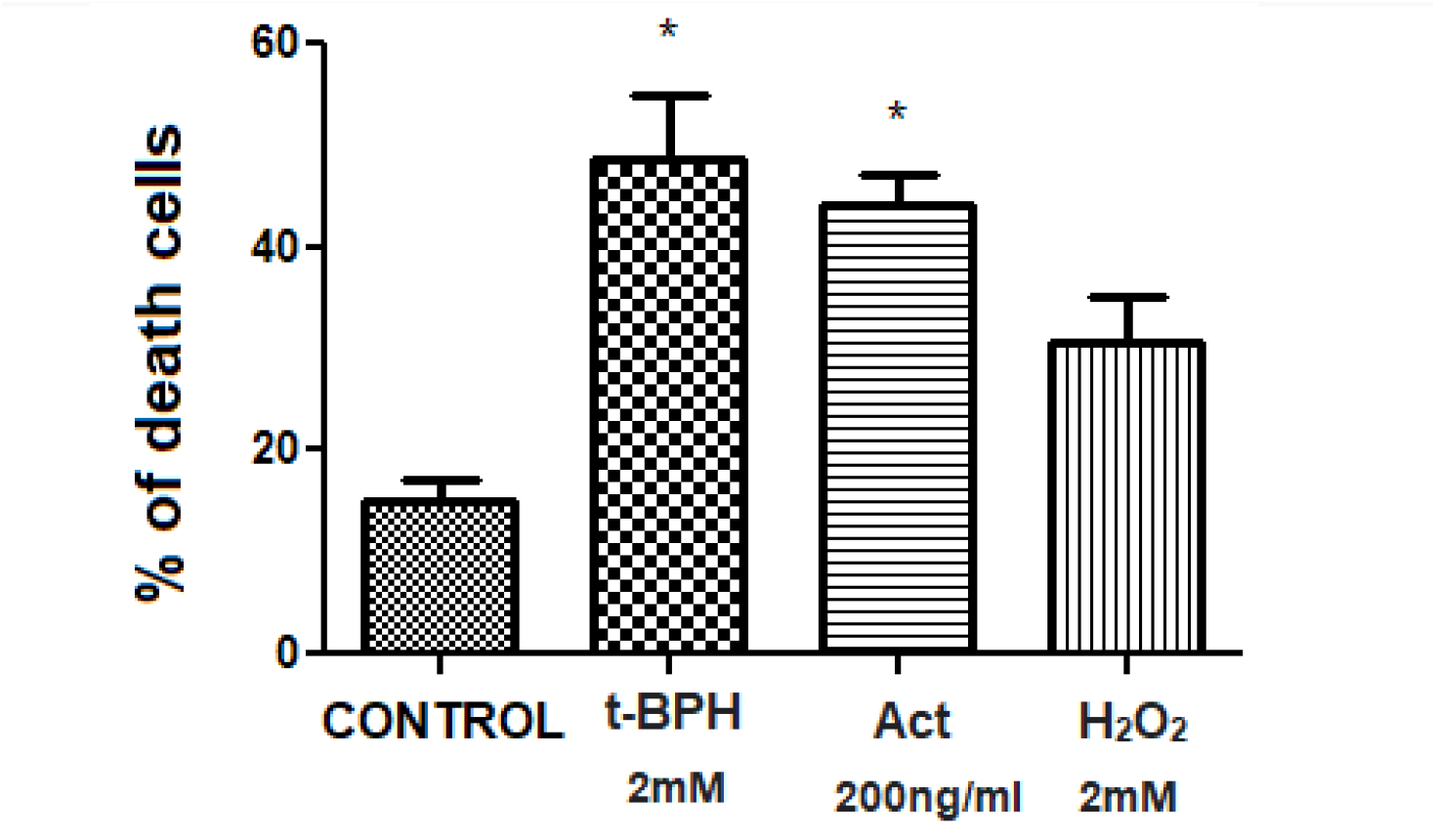
SF-9 cells treated with or without the apoptosis inducer t-BPH (2mM), Actinomycin D (200ng/ml), or H2O2 (2mM). After 18 h of exposure, the cells were stained with trypan blue and counted using a Newbauer chamber. *p<0.05, compared to the control (ANOVA and Newman-Keuls multiple comparisons).

### 4.5 Anti-apoptotic action of crude hemolymph from Podalia sp

The protection of hemolymph from ***Podalia sp***. (2% v/v) on **Vero** cells treated with t-BPH (0,5 or 1 mM) after 18 h was determined. Cell viability was determined using the **MTT** assay. As can be seen, the hemolymph of *Podalia sp* reduces the induction of apoptosis death by t-BPH (**figure 10, 11**).

**Figure 10:**
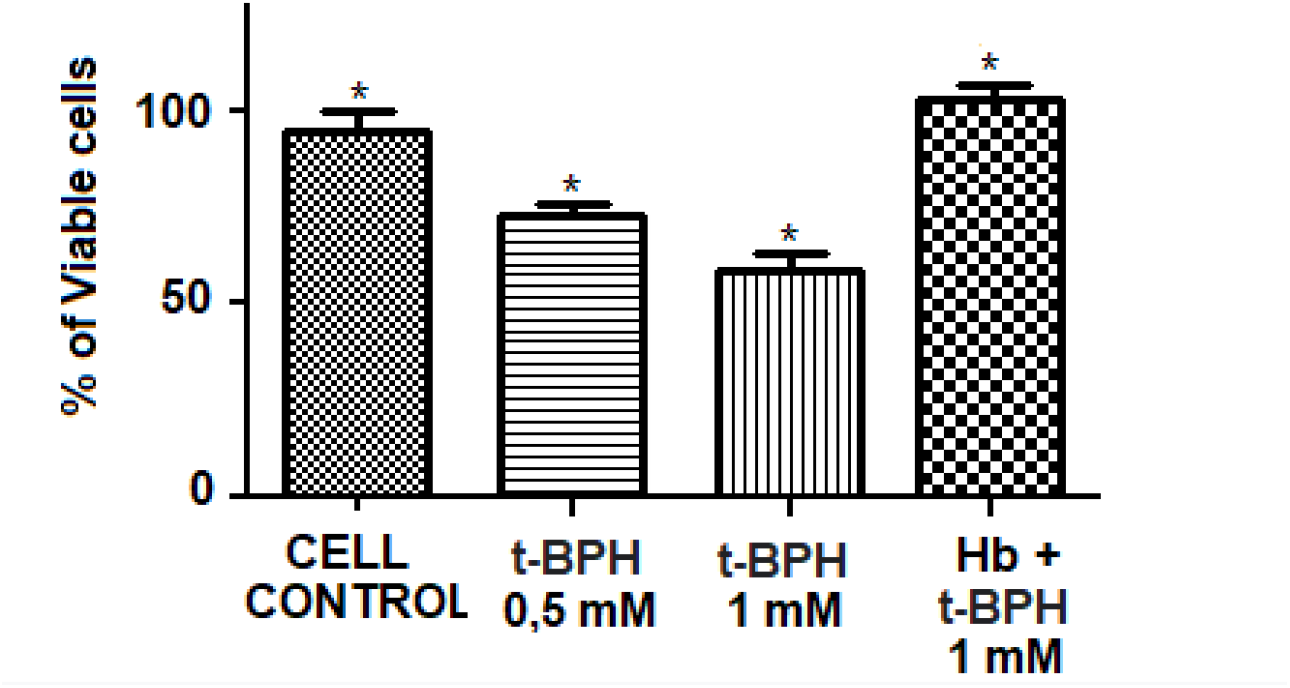
Viability of **Vero cells** (by MTT) after 18 hours of exposure with **t-BPH** (1mM) and treated with crude hemolymph from ***Podalia sp*** (2% v/v)

**Figure 11:**
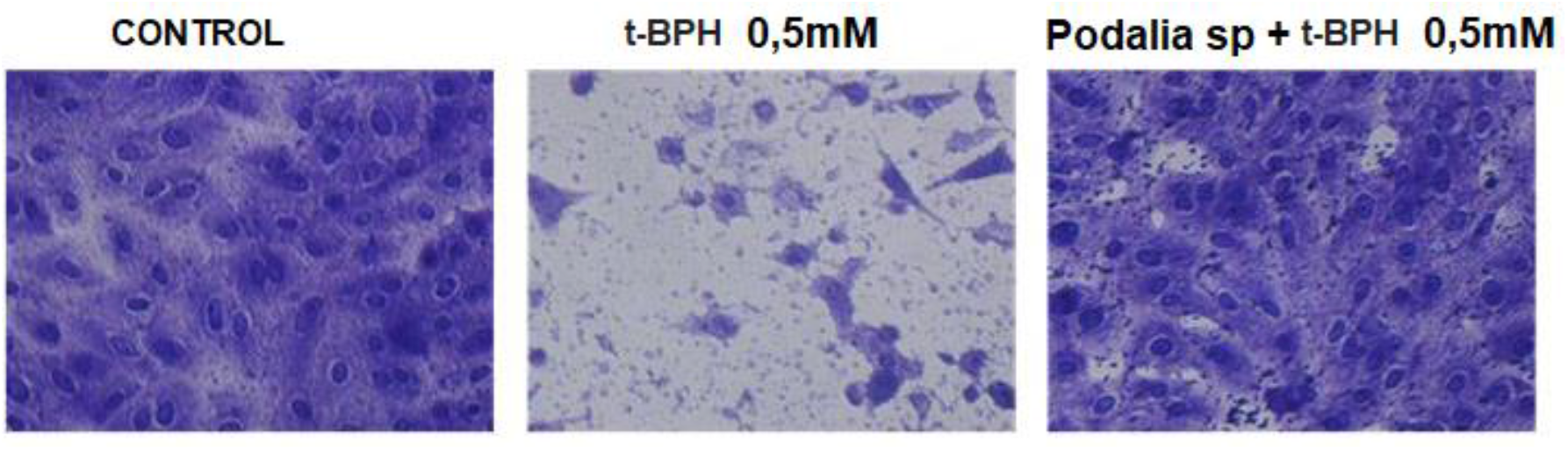
Effect of *Podalia sp*. hemolymph on protection against cell death induced by chemical inducers of apoptosis. VERO cells were pre-treated with or without 2% (v/v) crude hemolymph for 1 h. After this period, cell death was induced with 0,5mM of t-BPH for 18 hours. Cells were stained with crystal violet and observed under an inverted optical microscope (400x).

To verify whether the hemolymph of ***Podalia sp***. promotes cellular protection against **H**_**2**_**O**_**2**_, **VERO** cells were treated with 1% v/v of the hemolymph of *Podalia sp*. 1 h before the addition of H_2_O_2_ (25, 50 or 100μM), (figure 12). As can be seen, total hemolymph from *Podalia sp* was able to protect VERO cells from death induced by H_2_O_2_

**Figure 12:**
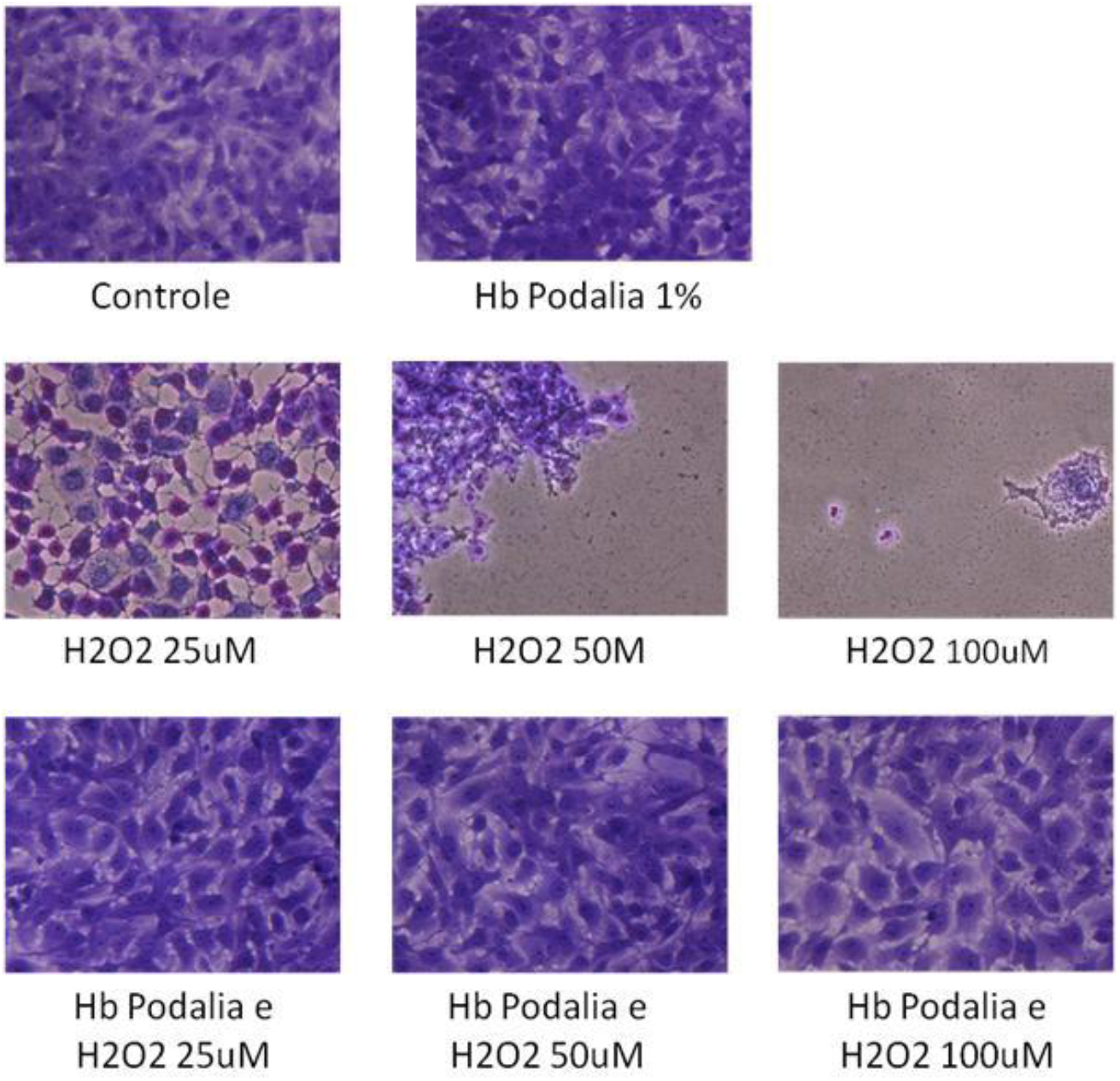
Effect of hemolymph in protecting cell death induced by Hydrogen Peroxide (H_2_O_2_). VERO cells were pre-treated with 1% (v/v) of total hemolymph from Podalia sp. and after 1-hour cell death was induced with H_2_O_2_, at concentrations of 25, 50 or 100μM for 18 h. The cells were stained with crystal violet and observed under an inverted optical microscope at 400x magnification.

### 4.6 Anti-apoptotic action of crude hemolymph from *M. albicolis*

To study the anti-apoptotic effect of *M. albicolis* hemolymph, **VERO** cells were pre-treated with or without total hemolymph (1% v/v) for 1 h. After this period, cell death was induced by H_2_O_2_. (0,4mM) (**figure 13a**) or t-BPH (0,5mM) (**figure 13b**) for 18 h. The total hemolymph from *M. albicolis* neutralized the effects of apoptosis inducers.

**Figure 13a:**
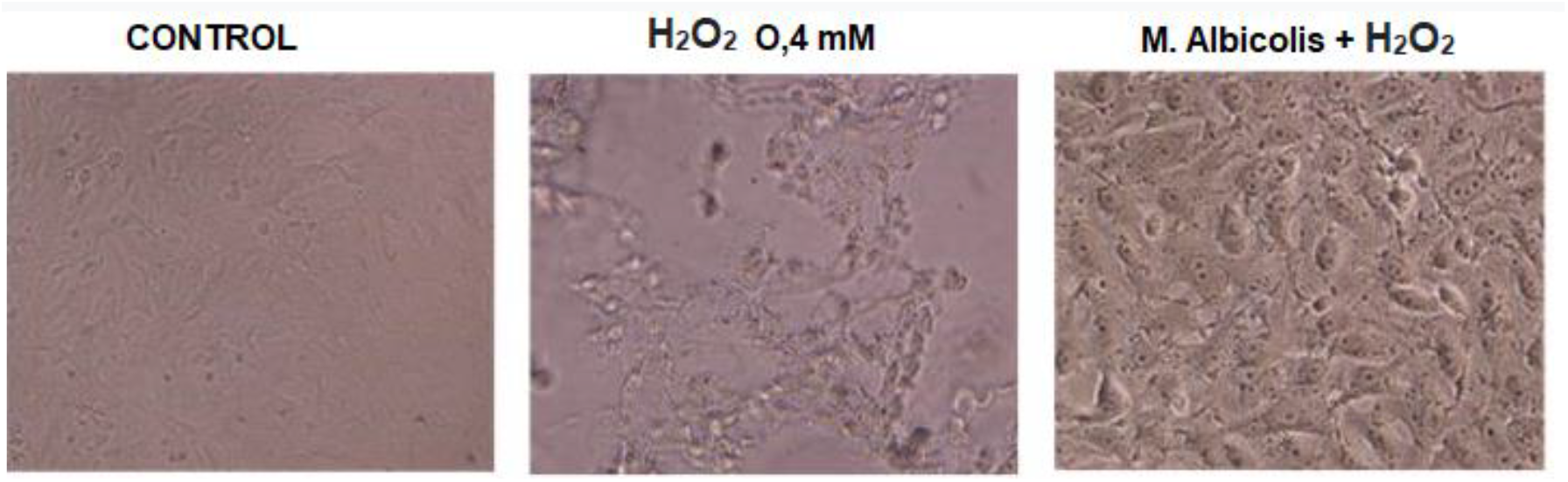
Effect of ***M. albicolis*** hemolymph on protection against cell death induced by chemical inducers of apoptosis. VERO cells were pre-treated or not with 1% (v/v) crude hemolymph for 1 hour. After this period, cell death was induced with 0,4 mM of H_2_O_2_. for 18 hours. The cells were observed in an inverted optical microscope (400x).

**Figure 13b:**
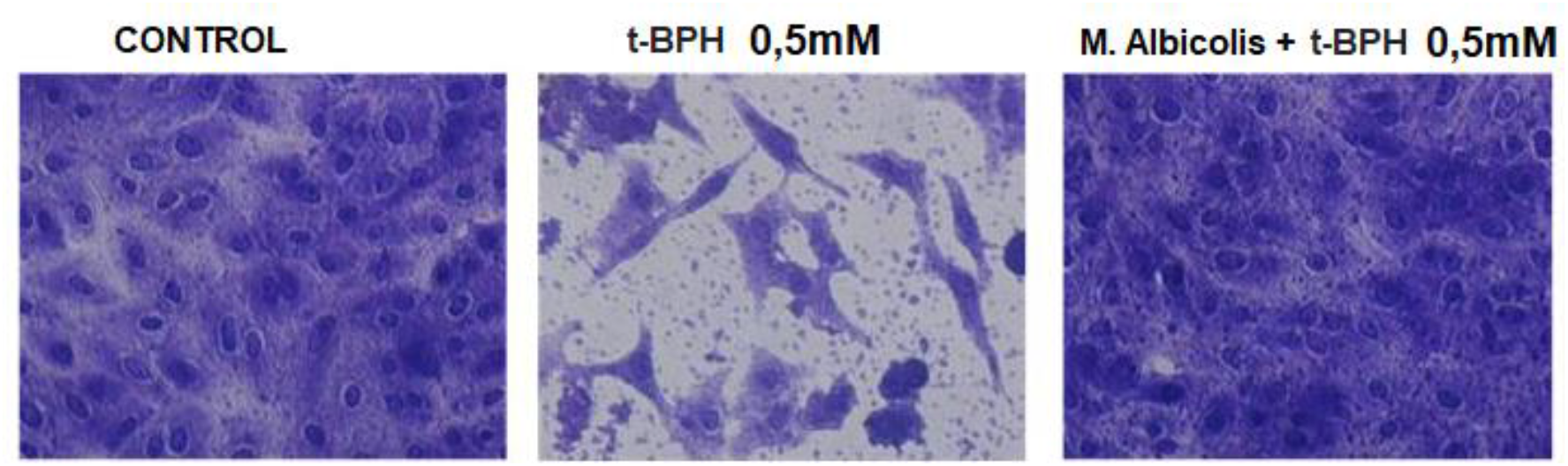
Effect of *M. albicolis* hemolymph on protection against cell death induced by chemical inducers of apoptosis (t-BPH). VERO cells were pre-treated with 1% (v/v) crude hemolymph for 1 h. After this period, cell death was induced with 0,5 mM of t −BPH for 18 hours. Cells were stained with crystal violet and observed under an inverted optical microscope (400x).

### 4.7 Analysis of the anti-apoptotic effect of hemolymph from *Podalia sp*. and *M. albicolis* by staining acridine orange (AO) and ethidium bromide (BE) with AO/BE

Cell death by Apoptosis in **SF-9** cell cultures was also determined by confocal microscopy after staining with acridine orange (AO) and ethidium bromide (BE) fluorophores. For this, SF-9 cells were treated with 1% v/v of hemolymph or its fractions 1 h before the addition of (t-BPH) at a concentration of 1mM (**Figure 14 a**) or Actinomycin D at a concentration of 800ng/mL (**Figure 14 b**). The total hemolymph of *Podalia sp*. and *M. albicolis* showed anti-apoptotic effects with both inducers. Viable cells stained uniformly green, whereas apoptotic cells stained red.

**Figure 14 a:**
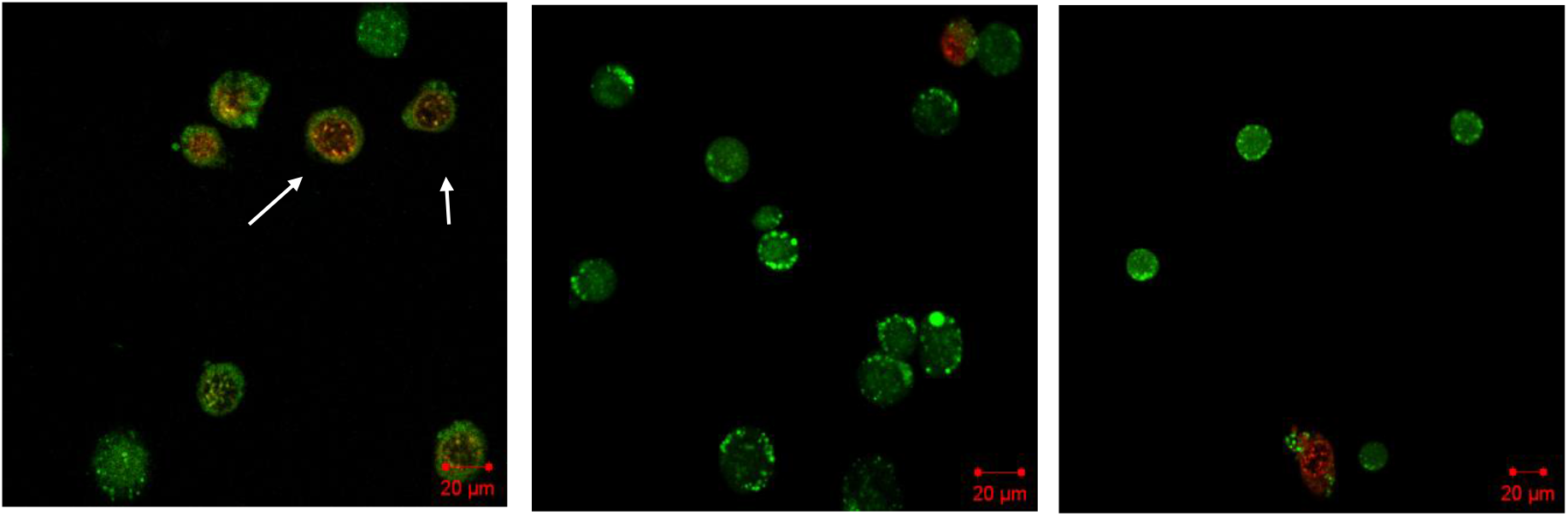
Morphology of SF-9 cells stained with Acridine Orange and Ethidium Bromide. A) cells treated with **1mM t-BPH**; B) cells treated with 1mM t-BPH and *M. albicolis* hemolymph; C) cells treated with 1mM t-BPH and *Podalia sp* hemolymph (arrow: apoptotic cells).

**Figure 14 b:**
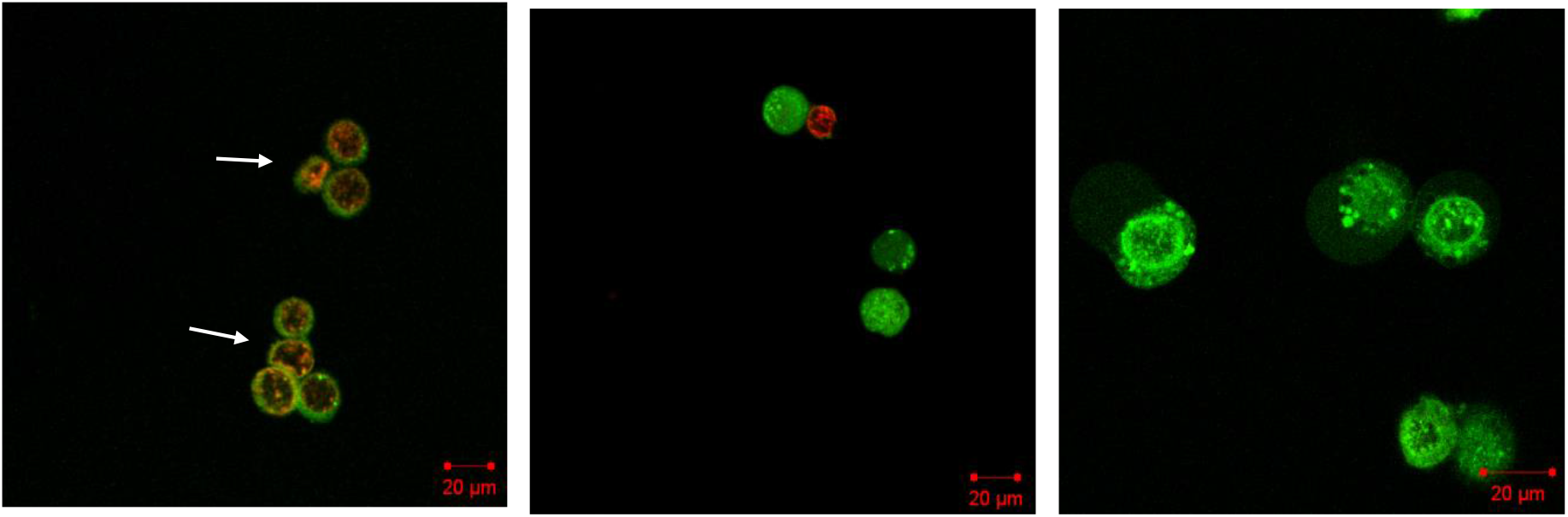
Morphology of SF-9 cells stained with Acridine Orange and Ethidium Bromide. A) cells treated with Actinomycin D 800ng/ml; B) cells treated with Actinomycin D 800ng/mL and *M. albicolis* hemolymph; C) cells treated with Actinomycin D 800ng/mL and *Podalia sp* hemolymph (arrow: apoptotic cells).

VERO cells also were maintained in PBS for 24 h or in PBS plus 1% v/v of total hemolymph from *Podalia sp*. As can be observed above, hemolymph protected VERO cells from death induced by nutrient depletion (**figure 15**).

**Figure 15:**
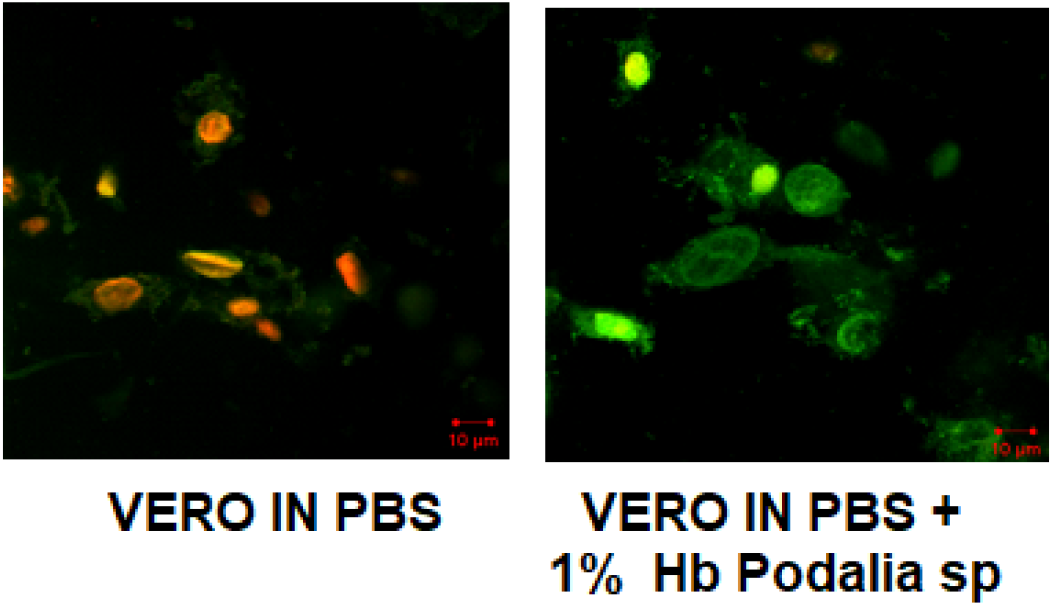
Effect of hemolymph on protection from cell death induced by nutrient depletion. VERO cells were kept in PBS for 24 h and treated with 1% (v/v) total hemolymph from *Podalia sp*. The cells were stained with acridine orange and ethidium bromide and observed under a confocal microscope.

### 4.8 Analysis of the anti-apoptotic effect of Podalia hemolymph in flow cytometry

The determination of DNA content by flow cytometry has been widely used to identify apoptosis in cell cultures. For this, **VERO** cells were treated with 1% v/v *Podalia sp*. hemolymph 1 h before adding t-BPH at a concentration of 0,5mM. **Figure 16** shows a histogram of the control culture, with a typical DNA profile of these cells, where the G1, S, and G2 phases are visible, and the sub-G1 population is low. In cultures where cells were treated only with t-BPH, an increase in the sub-G1 population was observed, indicating a process of death by apoptosis. In the culture where cell death was induced, but with the previous addition of hemolymph, there was an increase in populations in other phases of the cycle, indicating that hemolymph acts by stimulating cell proliferation and preventing cell death by apoptosis

**Figure 16:**
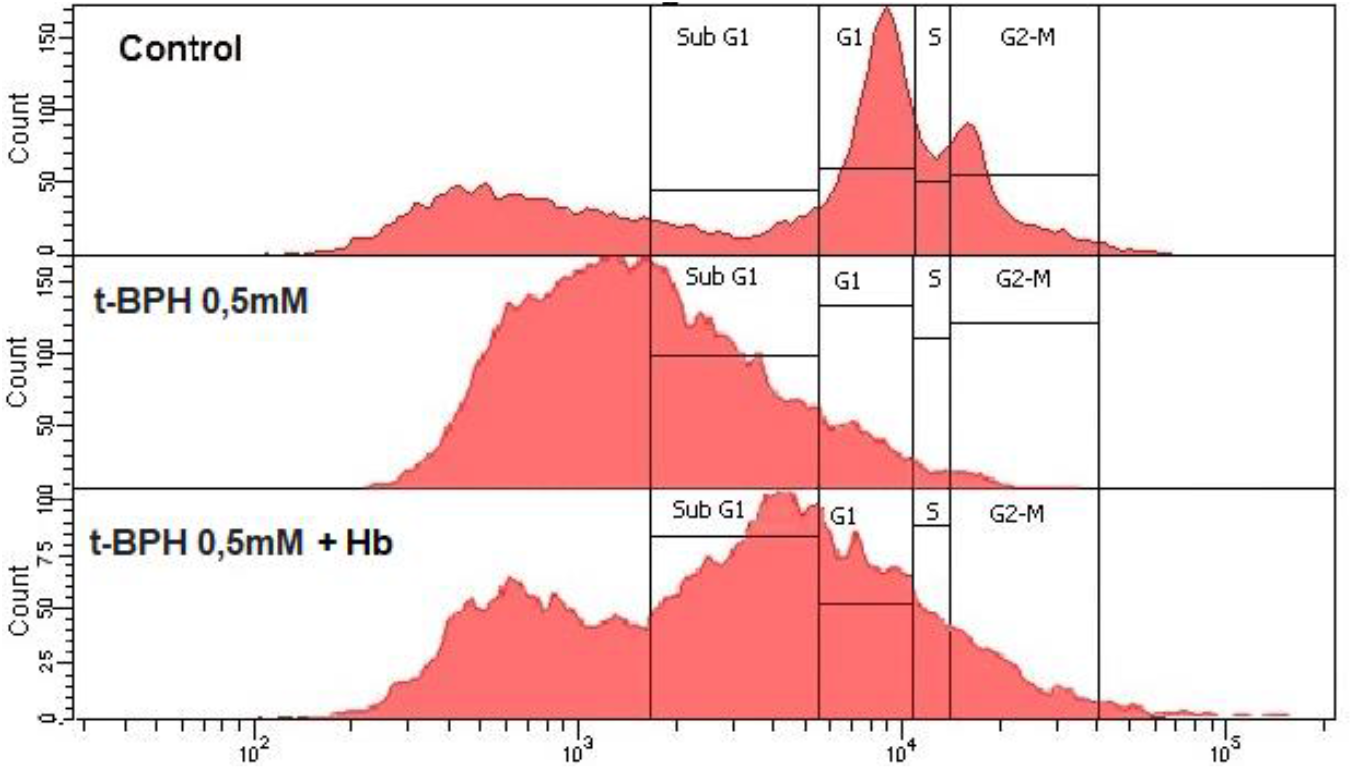
Histogram of the DNA of VERO cells treated with 1% v/v of *Podalia sp* hemolymph (Hb), 1 hour before the addition of t-BPH at a concentration of 0,5 mM.

### 4.9 Determination of alteration of the cytoskeleton of VERO cells by labeling with phalloidin-FITC

Studies have provided important information about how the actin cytoskeleton contributes to the control of growth, both in transformed and non-transformed cells, indicating its participation in the regulation of several cellular processes related to transformation, such as proliferation, anchorage-independent growth, and contact inhibition of growth (Pawlak and Helfman, 2001). Cellular transformation into fibroblasts, as well as the process of apoptosis, is characterized by several morphological changes, with a rounded phenotype, a decrease in actin bundles of the cytoskeleton, and striking features of stress fibers (Janmey and Chaponnier, 1995). In control cells labeled with phalloidin-FITC, a network of highly organized actin filaments was observed, with abundant stress fibers, most of which had an orientation parallel to the major cell axis. With the induction of cell death by t-BPH, there was a significant reduction in the number of stress fibers and shortening of those that remained. To study the possible effect of hemolymph in maintaining the cytoskeleton of cells induced apoptosis, VERO cells were pre-treated with *Podalia sp*. hemolymph and treated with 100μM μM t-BPH for 4 h. When VERO cells were pre-treated with total hemolymph from *Podalia sp*., before apoptosis induction, the cytoskeleton was maintained, showing the stress fibers clearly evident. Therefore, the hemolymph of *Podalia sp*. showed a protective effect by blocking or mitigating disruption of the cytoskeleton **(Figure 17)**.

**Figure 17:**
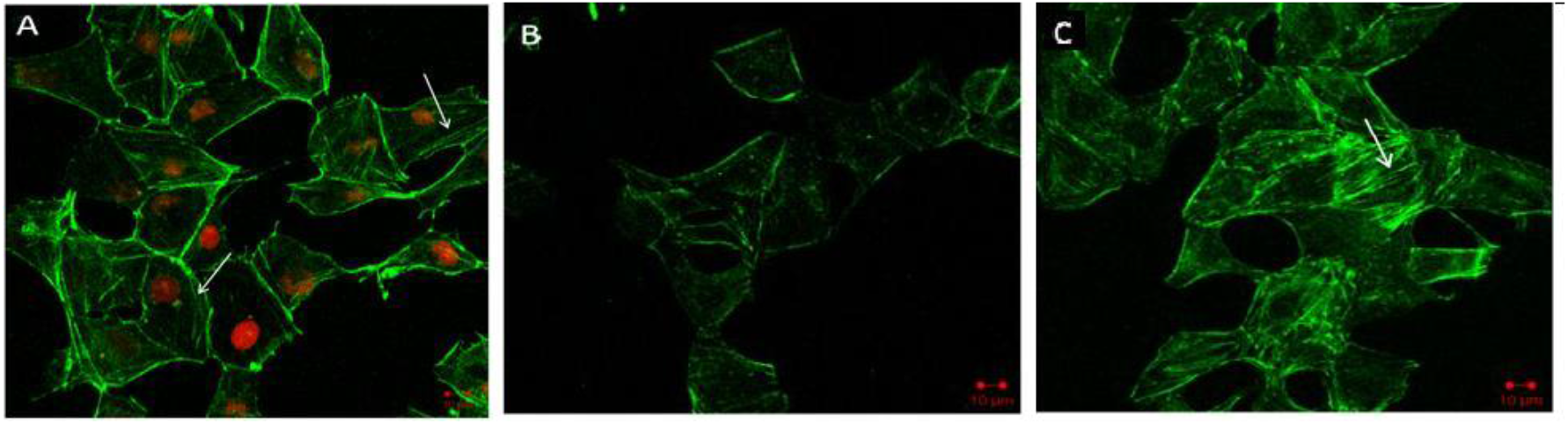
Actin filaments in Vero cells (phalloidin labeling). (A) Control VERO cells showing abundant fibers parallel to the major cell axis (arrows). The nuclei were stained with propidium iodide. (B) VERO cells treated with t-BPH (100μM) for 4 h. (C) Cells were pretreated with total Podalia hemolymph (1% v/v) for 1 h. After this period, they were treated with t −BPH at a concentration of 100μM for 4 hours. In cells pretreated with hemolymph, the cytoskeleton was maintained, showing clearly evident stress fibers (arrow). The nuclei were not marked. The cells were then analyzed using confocal microscopy.

Likewise, VERO cells were pre-treated with hemolymph of *M. albicolis* total for 1 h before apoptosis induction with 100μM t-BPH. As can be seen, the cytoskeleton and nucleus were also maintained in the culture treated with M. albicolis hemolymph, demonstrating a protective effect, blocking or mitigating the disruption of the cytoskeleton **(figure 18)**.

**Figure 18:**
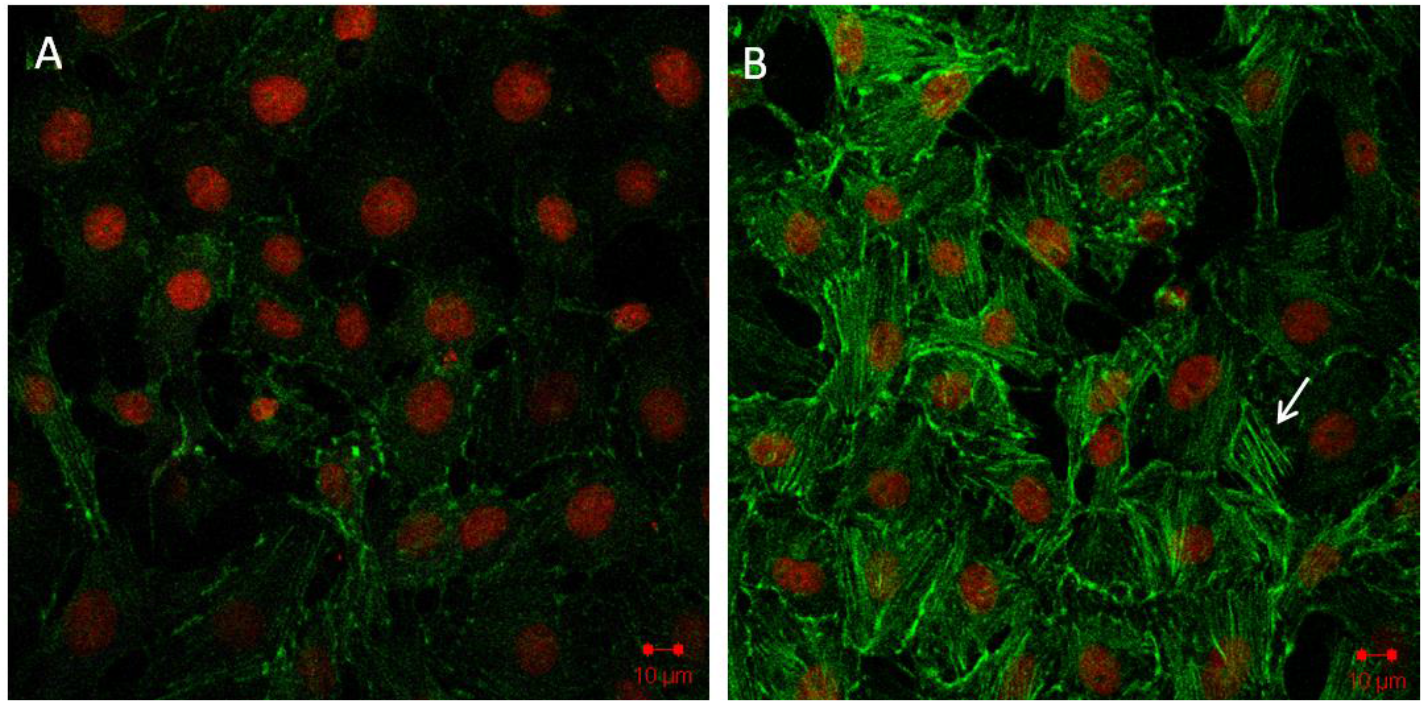
Protector effect of *M. albicolis* in actin filaments of Vero cells (phalloidin labeling). (A) Cells treated with t-BPH at a concentration of 100μM for 4 hours. (B) VERO cells pre-treated with *M. albicolis* total hemolymph (1% v/v) for 1 h. After this period, cells were induced to die with t-BPH at a concentration of 100μM, for 4 h. In cells pre-treated with hemolymph, the cytoskeleton was maintained, showing clearly evident stress fibers (arrow). The core was then labeled with propidium iodide. Cells were analyzed by confocal microscopy.

### 4.10 Labeling of tubulin in VERO cells

The addition of hemolymph also maintained the tubulin structure in VERO cells even after the addition of 100μM t-BPH. Microtubules were evidenced by a primary anti-β-tubulin antibody, followed by a secondary anti-mouse IgG antibody conjugated to Alexa Fluor 488 **(Figure 19)**. Microtubules were distributed throughout the cytoplasm of cells, similar to the organization of normal cells. Mitotic cells with microtubule organization characteristics of normal mitotic spindles were also observed.

**Figure 19:**
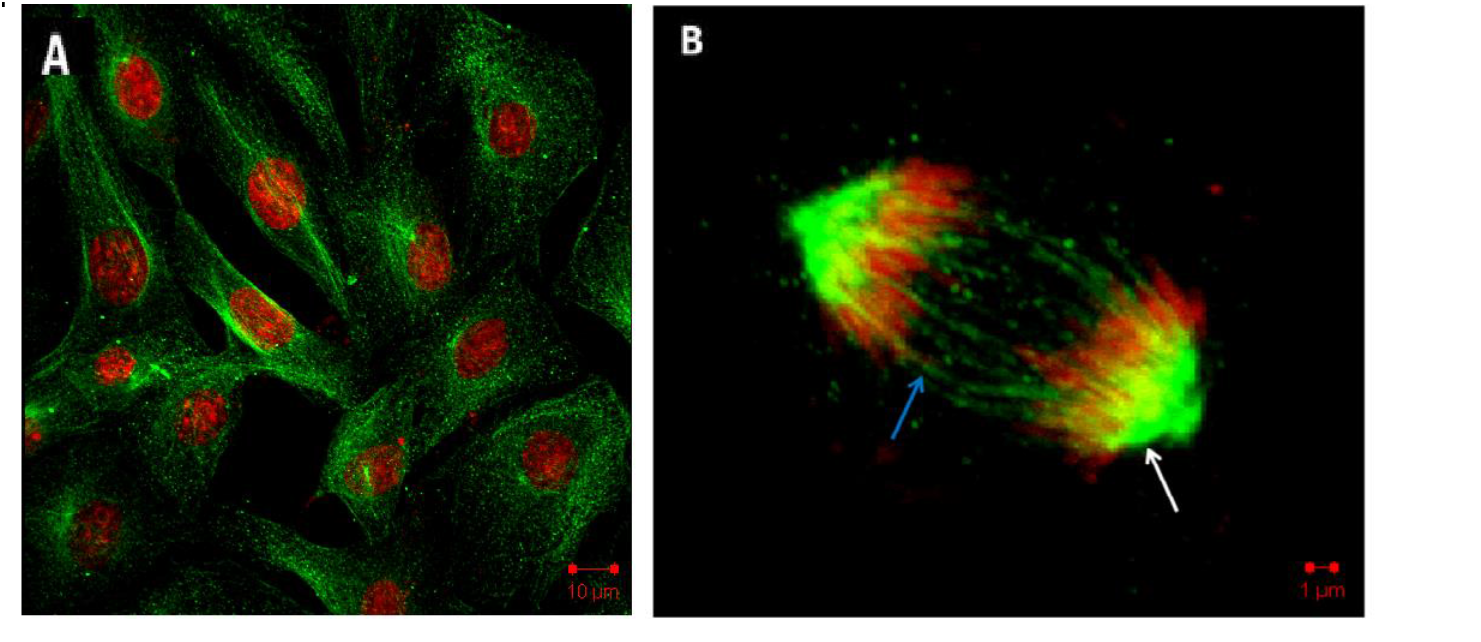
Microtubules in Vero cells after induction of apoptosis with 100μM t-BPH but pre-treated with M. albicolis total hemolymph (1% v/v) for 1 hour. Microtubules were evidenced by primary anti-β tubulin antibody, followed by secondary anti-mouse IgG antibody conjugated to ALexa Fluor 488. The nucleus was labeled with propidium iodide. Chromosomes are highlighted in red and microtubules in green. The blue arrow points to polar microtubules and the white arrow points to astral microtubules. Cells were analyzed by confocal microscopy.

### 4.11 Determination of mitochondrial membrane potential (ΔΨm) in SF-9 cells

Oxidative stress is related to the loss of mitochondrial integrity and triggers the intrinsic pathway of apoptosis. We used the dye JC-1 (5,5’,6,6’-tetrachloro-1,1’,3,3’-tetraethylbenzimidazolylcarbocyanine iodide) to evaluate the integrity of the mitochondrial membrane of **Sf-9** cells pretreated with or without *Podalia sp* hemolymph and induced apoptosis with t-BPH (600μM) for 4 hours. Red staining indicates healthy mitochondria with preserved membrane potential (A, C). Green staining indicates the loss of mitochondrial membrane electrical potential (B). (Figure 20). The **figure 20** show representative images of JC-1 staining using confocal microscopy.

**Figure 20:**
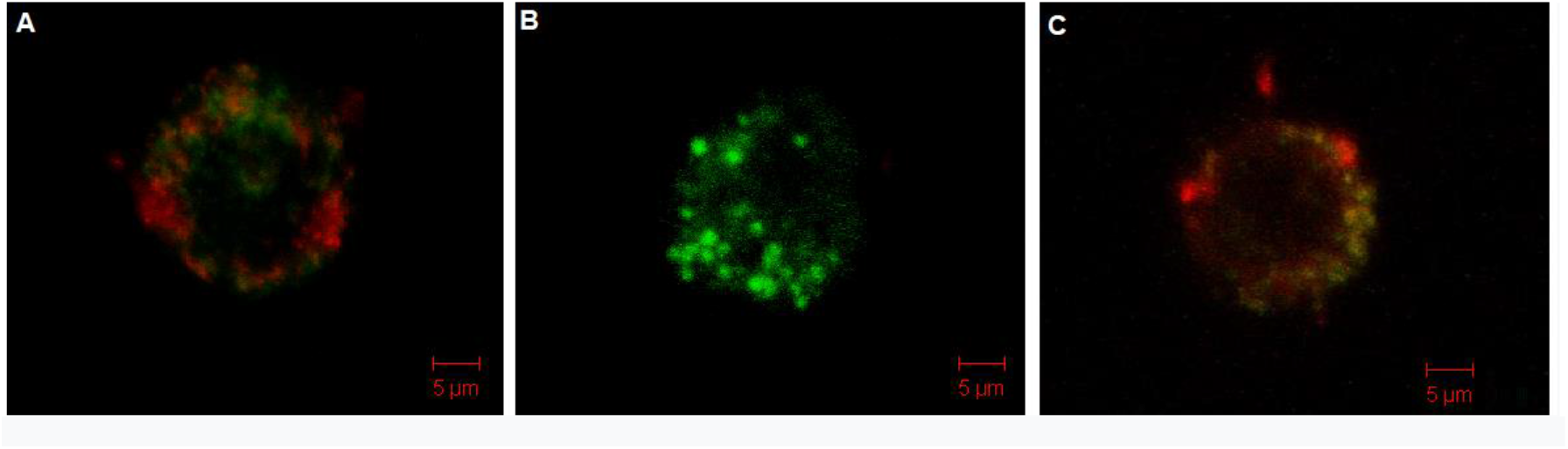
Sf-9 cells labeled with JC-1. (A) control cells; (B) cells treated with t-BPH (600μM) for 4 hours; (C) Cells pre-treated with hemolymph OF *Podalia’s* total (1 hr) and treated with t-BPH (600μM) for 4 hours.

Mitochondria are involved in many processes that are essential for cell survival, including energy production, redox control, calcium homeostasis, and certain metabolic and biosynthetic pathways. In addition, mitochondria play an essential role in the cell death process known as apoptosis. This mitochondrial pathway of death by apoptosis can cause various diseases such as cancer, diabetes, ischemia, and neurodegenerative disorders such as Parkinson’s and Alzheimer’s diseases (Bouchier-Hayes et al., 2005). Owing to the numerous activation mechanisms of death by apoptosis, the determination of the mechanism of action of different anti-apoptotic proteins, especially those with a broad spectrum of action, whether inhibiting death by different inducers or preventing this death in different cell types, can be of great importance. High medical and biotechnological interest. Studies with an anti-apoptotic protein isolated from the hemolymph of Lonomia obliqua showed that its likely action is on the mitochondrial membrane, maintaining a high electrical potential and preventing membrane permeability (Mendonça & Martins, 2022).

The hemolymph of *Podalia sp* and M. albicolis showed a very similar action as the anti-apoptotic protein of Lonomia obliqua. It is also suggested that the site of action of this protein could be similar to that of the Bcl-(x) family proteins, since, in all experiments, a block or decrease in death by apoptosis was observed, with evident protection of the potential of the mitochondrial membrane. Due to the broad spectrum of action of hemolymph, which acts both in insect and mammalian cells, either by inducing apoptosis by chemical or nutritional agents, there is a molecule present in the hemolymph that is capable of acting as a general suppressor of the apoptosis process, for example, a conserved pathway in many organisms.

The addition of hemolymph to cultures not only neutralizes or reduces chemically induced death, but also physiological death that normally occurs due to culture stress.Cells treated with hemolymph remained alive for longer, indicating that this protein could also be used as a supplement to culture media. By blocking cell death, the cell remains functional, which is a fundamental indication for the use of a molecule with these characteristics as a drug against diseases neurodegenerative as Parkinson’s and Alzheimer’s diseases.

The hemolymph of the caterpillars *Podalia sp*. and *M. albicolis* showed an anti-apoptotic effect both in insect cells and in mammalian cells, indicating that this protein eventually acts as a basic mechanism of cell death.

Silkworm hemolymph has been reported to inhibit apoptosis in the later stages of culture, both in insects and human cells (Choi et al., 2002). Studies have shown the occurrence of cell death by apoptosis in cultures of mammalian and insect cells with nutrient depletion (Mendonca et al., 2002; Meneses-Acosta et al., 2001) or by the induction of chemical agents (Souza et al., 2005). Cell death is a very important factor in production processes that limits the industrial production of proteins of economic interest. One way to increase cellular productivity is to inhibit or attenuate cell death. We have previously demonstrated the presence of a potent anti-apoptotic protein in the hemolymph of Lonomia obliqua, which extends cell culture viability through apoptosis prevention (Maranga et al., 2003; Raffoul et al., 2005; Souza et al., 2005). Has been reported also that mitochondria has one important action in the apoptosis control process, being that Mitochondria Membrane Permeabilization (MMP) an important stage in this process. We have also shown that the addition of Lonomia obliqua hemolymph in the culture leads to a prolongation of the cell, leading to a high electrochemical potential of the mitochondria, avoiding the loss of membrane permeability and cytochrome Cytochrome-C release. Thus, even though they originate from different species, we believe that the hemolymph of *Podalia sp*. and *M. albicolis* can act through similar mechanisms to the hemolymph of Lonomia obliqua. However, the hemolymph from *Podalia sp*. And *M. albicolis* caterpillars did not prove to be cytotoxic or genotoxic to cell cultures at concentrations below 5%, which may be useful as a cell cultivation additive. Even though the electrophoretic profiles of both caterpillars were different, the effects were similar in both mammalian and insect cells, which may indicate the presence of components with similar functions and mechanisms of action. Crude hemolymph from both caterpillars showed anti-apoptotic effects in VERO and SF-9 cells pre-treated with only 1% v/v of hemolymph and induced death by different, potent apoptotic inductors such as terbutyl, actinomycin D, hydrogen peroxide, and depletion of nutrients; this protective effect blocked and attenuated the disruption of the cytoskeleton (actin filaments); this protective effect was also observed on the integrity of the mitochondrial membrane of SF-9 cells pre-treated with both hemolymph and treated with the apoptosis inducer terbutil at concentrations of 25–100μM.

## 5. CONCLUSION

The hemolymph of *Podalia sp*. and *M. albicolis* caterpillars was neither cytotoxic nor genotoxic to cell cultures at concentrations below 5%.

The electrophoretic profile of the hemolymph of *Podalia sp*. was different from that of *M. Albicolis*.

This was observed in the hemolymph of *Podalia sp*. caterpillars and *M. albicolis* molecules of pharmacological and biotechnological interest.

Crude hemolymph from both caterpillars showed anti-apoptotic effects in both VERO and SF-9 cells, against different inducers such as terbutyl, actinomycin D, and hydrogen peroxide, and by depletion of nutrients.

The hemolymph of both caterpillars showed a protective effect, blocking or attenuating the disruption of the cytoskeleton (actin filaments) of VERO cells treated with the apoptosis inducer terbutil at concentrations of 25–100μM.

Protective effect was observed on the integrity of the mitochondrial membrane of SF-9 cells pre-treated with both hemolymphs and treated with the apoptosis inducer Terbutil at concentrations of 25 to 100μM

By acting on the mitochondrial pathway of death by apoptosis, a pathway that can cause disorders and diseases neurodegenerative such as Parkinson’s and Alzheimer’s diseases, substances present in the hemolymph of these caterpillars could be good candidates in studies for the treatment of these diseases.

## Abbreviations

HB: Hemolymph;
t-BPH: Tert-Butylhydroperoxide;
PI: Propidium Iodide;
AO: acridine orange;
EB: ethidium bromide;
(DiOC6 (3): 3,3’Dihexyloxacarbocyanine Iodide;
JC-1: 3,3’E-Tetraethylbenzimidazolylcar-bocyanine Iodide;
MMP: Mitochondrial Membrane Permeabilization.

## Acknowledgments

The authors acknowledge the financial support received from FAPESP:2010/52434-6 and a doctorate fellowship to Nathalia Delazeri de Carvalho from FAPESP-2011/50095-2

